# Exploring Warming Effects on lower food-web dynamics in the plankton of the River Elbe Estuarine Ecosystem in summer: Insights from a Mesocosm Experiment

**DOI:** 10.64898/2026.04.09.714476

**Authors:** Luisa Listmann, Julia Golebiowska, Max Lambrecht, Sahed Ahmed Palash, Diana Nicole Puerto Rueda, Hans-Peter Grossart, Arne M. Malzahn, Elisa Schaum, Nicole Aberle

**Affiliations:** Institute of Marine Ecosystem and Fisheries Science, University of Hamburg, Hamburg, Germany; Leibniz Institute of Freshwater Ecology and Inland Fisheries (IGB), Dept. Plankton and Microbial Ecology, Zur alten Fischerhuette 2, S-16775 Stechlin, Germany; Institute of Biochemistry and Biology, Potsdam University, Maulbeerallee 2, D-14469 Potsdam, Germany (second affiliation HPG); Department of Microbiology and Biotechnology, Institute for Plant Sciences and Microbiology University of Hamburg, Ohnhorststraße 18, Hamburg 22609, Germany

**Keywords:** plankton community, mesocosm, warming, Elbe river

## Abstract

Understanding how warming alters estuarine plankton communities is essential for predicting future changes in biodiversity and ecosystem functioning. We conducted a four-week indoor mesocosm experiment using natural summer plankton from the Elbe River to examine the effects of warming (+2 °C and +4 °C) on abiotic conditions and responses of the plankton community. In this study, oxygen concentrations, primary producer biomass (chlorophyll a, microphytoplankton) and microzooplankton abundances declined sharply during the first 10 days across all treatments while mesozooplankton abundances increased. This suggests a strong top-down control by mesozooplankton on lower trophic levels across all temperature treatments. Primary producers’ biomass and oxygen concentrations recovered after an initial decline, however to lower levels compared to the onset of the experiment while micro- and mesozooplankton remained low during the second half of the experiment. Nutrient dynamics indicated progressive remineralization, with increasing ammonium, NOx, and silicate concentrations, while phosphate concentrations remained low throughout the experiment. Complementary DNA and RNA metabarcoding revealed similar community turnover over time in all treatments and temperature effects became only pronounced at the end of the experiment. Overall, warming effects were subtle relative to the strong internal trophic dynamics likely caused by the artificial mesocosm setup. Our findings of changes in plankton community dynamics indicate that biotic interactions, changes in trophic diversity and other environmental factors, i.e. oxygen concentrations are likely the drivers of this estuarine system rather than warming alone.

## Introduction

Global warming is one of the most pervasive drivers of change in aquatic ecosystems, profoundly influencing biodiversity, community composition, and ecosystem functioning across freshwater, estuarine, and marine environments. Rising temperatures directly affect physiological rates of organisms, altering growth, metabolism, and phenology while indirectly reshaping species interactions and trophic pathways (Hillebrand et al. 2018; Yu et al. 2025). In planktonic systems, these changes are particularly pronounced at the base of the food web, where bacterioplankton, phytoplankton, and microzooplankton regulate primary production, nutrient cycling, and the transfer of energy to higher trophic levels including mesozooplankton (Finkel et al. 2010; Sarker et al. 2020). Warming has been shown to modify plankton community, with implications on elemental stoichiometry and nutrient fluxes, and to enhance the relative role of microbial pathways in carbon cycling (Šorf et al. 2014; Finkel et al. 2010; Cherabier and Ferrière 2022). Such temperature-driven shifts in lower food-web dynamics can cascade through aquatic ecosystems, ultimately influencing productivity, stability, and resilience under ongoing climate change.

Phytoplankton blooms are major, recurrent phenomena that play a crucial role in fueling secondary production in aquatic ecosystems (Sommer et al. 2011; Edwards and Richardson 2004). Under ongoing climate change, pronounced shifts in bloom phenology and community composition have been widely documented. These include advances in the timing of the spring phytoplankton bloom (Winder et al. 2012; Weyhenmeyer, Blenckner, and Pettersson 1999) as well as changes in taxonomic and size structure (Sommer et al. 2007a; Daufresne, Lengfellner, and Sommer 2009). Such alterations can substantially affect primary production in lotic and estuarine systems, where enhanced river runoff and nutrient loading further modify phytoplankton dynamics (Savoy et al. 2019). The formation of blooms is dependent on the growth rate of primary producers, i.e. exceeding their losses. These losses include grazing by micro- and mesozooplankton, mortality or loss due to perturbation and mixing (Irigoien, Flynn, and Harris 2005). Further, different grazers, such as copepods or cladocerans, might modulate the size structure and community composition of phytoplankton blooms (Sommer, Hansen, and Sommer 2006). In addition, this balance is influenced by the differential temperature dependencies of autotrophic (i.e. primary producers) and heterotrophic processes (i.e. micro- and mesozooplankton) since photosynthesis is considered to be primarily limited by light, while heterotrophic metabolism is highly temperature-dependent (Metabolic Theory of Ecology) (Brown et al. 2004).

Microzooplankton (MicroZP) in the size range of 20-200 µm, comprising mainly of eukaryotic protozoa and small metazoans, is highly sensitive to temperature fluctuations (Aberle et al. 2012) and constitutes a pivotal component of the microbial food web. Increased temperatures can accelerate MicroZP growth and grazing and alter the dynamics of phytoplankton communities and bloom phenology (Aberle et al. 2012; Wohlers et al. 2009; Sarmento et al. 2010). MicroZP is regarded as the principal grazer of phytoplankton (Landry and Calbet 2004; Aberle, Lengfellner, and Sommer 2007), holding the potential to modulate and suppress phytoplankton standing stocks (Sherr and Sherr 2009; Löder et al. 2011). Alterations in growth and grazing by MicroZP, alongside changes in net primary production and phytoplankton community composition under warming (Schaum et al. 2017), can have profound effects on the timing, magnitude, community composition, and diversity of phytoplankton, significantly impacting estuarine food web functioning and carbon transfer efficiencies (Isla, Lengfellner, and Sommer 2008; Aberle et al. 2012).

Mesozooplankton (MesoZP) are important grazers in planktonic food webs (0.2-2 mm) relying on primary and secondary production of phyto- and microzooplankton (Klauschies et al. 2012; Aberle et al. 2015). In turn, MesoZP serve as prey for e.g. early live stages of fish, and adult planktivorous fish and represent the link to higher tropic levels (Aberle and Malzahn 2007). Increasing temperatures affect their metabolism directly (Isla, Lengfellner, and Sommer 2008), while alteration in nutrient loads affect mesozooplankton prey communities and their biochemical composition indirectly via lower trophic levels (Malzahn et al. 2007; Boersma et al. 2008).

The above-mentioned plankton community components can be measured using “classical” experimental approaches such as microscopy and flow cytometry (Holland et al. 2025). In addition, genomic data such as metabarcoding provide a novel tool to complement traditional assessments as they can reveal underlying biological mechanisms driving observed ecological responses and improve our ability to predict ecosystem change (Grossart et al. 2026). Genomic and metabarcoding approaches have been shown to capture shifts in plankton community composition and functional diversity that are often missed by microscopy alone, offering broader taxonomic resolution and insights into lower-trophic dynamics under environmental stressors (e.g., DNA metabarcoding in plankton warming and salinity studies; microscopy vs. genomic methods comparison) (Hall et al. 2025; Martens et al. 2024).

The River Elbe estuary is a highly productive and strongly modified aquatic system that exhibits pronounced seasonal variability in plankton dynamics and oxygen conditions. During summer, elevated temperatures and sustained nutrient availability frequently promote intense phytoplankton blooms, which are often followed by rapid biomass decay and enhanced microbial respiration. This tight coupling between phytoplankton production and heterotrophic degradation has been identified as a key driver of recurrent oxygen depletion in the freshwater and mesohaline reaches of the

Elbe, with substantial consequences for ecosystem functioning and biogeochemical cycling (Amann, Weiss, and Hartmann 2012; Schöl et al. 2014). Oxygen limitation in the Elbe is thus closely linked to lower food-web processes, where the ratio between primary production, grazing, and microbial decay determines carbon turnover and oxygen availability (García-Oliva et al. 2025; Steidle and Vennell 2024; Rabalais et al. 2010). However, investigating these mechanisms *in situ* is challenging due to strong hydrodynamic variability and longitudinal transport.

Mesocosm experiments offer a powerful methodological approach for investigating the intricate interplay of lower and higher trophic dynamics in estuarine ecosystems, particularly in the context of extreme weather events (i.e. extreme flooding and heat waves and see Grossart et al. 2026) and global warming (Hall et al. 2025; Sommer et al. 2007a). By simulating semi-natural environmental conditions in a controlled setting, mesocosms enable manipulations of specific variables such as temperature, nutrient levels, and salinity while closely monitoring the resulting biological responses (Vidussi et al. 2011; Trombetta et al. 2024; Stewart et al. 2013). This level of control allows for the isolation of variables that contribute to complex trophic interactions, providing insights into processes that are often difficult to discern in natural environments due to their inherent variability and short-term dynamics. Moreover, mesocosm experiments facilitate the examination of community dynamics over time, enabling the observation of phenological shifts and responses to warming in real-time (Aberle et al. 2012; Winder et al. 2012). We expect that experimental warming will shift the balance between autotrophic and heterotrophic processes in the Elbe plankton community by enhancing microzooplankton growth and grazing, leading to reduced phytoplankton biomass and altered bloom phenology. In addition, temperature-induced changes in plankton community composition and trophic interactions will increase microbial carbon turnover and respiration, resulting in enhanced oxygen consumption and reduced oxygen availability under warming conditions. These hypotheses were tested in the present study, which aims to elucidate the effects of warming on a summer plankton community from the River Elbe estuarine ecosystem and its associated abiotic environment. By doing so, this study enhances our understanding of the ecological implications of global change for aquatic systems and improves our ability to predict future changes in estuarine ecosystems.

## Methods

To analyse effects of warming on a freshwater riverine summer plankton community we conducted a four-week indoor mesocosm experiment from the 29^th^ of July to 23^rd^ of August 2024 at the Institute of Marine Ecosystem and Fisheries Science (IMF, Hamburg University) using natural river water from the Elbe taken from Bunthaus station (53.4618° N, 10.0645° E) (Martens, Biederbick, and Schaum 2024). The plankton community including viruses, bacteria, phytoplankton as well as micro- and mesozooplankton was sampled over four weeks (Table 1).

**Table 1:** Sampling and preservation of abiotic and biotic parameters during the experiment. All filters used were GF/F filters: NC45 membrane (cellulose nitrate).

| Sample | Volume | Processing | Preservation and storage | Data |
| --- | --- | --- | --- | --- |
| Abiotic parameters | NA | *in situ* |  | Data shown here |
| Nutrients | 3x 15 mL | Filtered on 0.45µm | Frozen at -80°C | Data shown here |
| Chlorophyll a | 150 mL | Filtered on 0.45 µm CF/F CN | Frozen at -80°C | Data shown here |
| Viruses and Bacteria | 1 ml | Filtered on 25µm | 1% Glutaraldehyde and storage at -80°C | Data not shown |
| Pico- and Nanoplankton | Direct measurement | Filtered on 25 µm |  | Data not shown |
| Microphytoplankton | 100 mL | No filtration | Neutral Lugol 0.5% and storage in the dark/temperature | Bulk abundances shown here |
| Microzooplankton | 200 mL | No filtration | Acidic Lugol 1% and storage in the dark/room temperature | Bulk abundances shown here |
| Mesozooplankton | 31.4 L | Net tow | 4% Formaldehyde and storage at room temperature | Bulk abundances shown here |
| rDNA/rRNA | 2 L | Filtration (see methods) | Filters were immediately frozen and stored in liquid nitrogen at -80°C | summarized metabarcoding data shown |

### Experimental Setup

The experimental set-up consisted of nine PVC tanks (1.2 m diameter, 1.0 m height, 860 L filling capacity, www.koi-ka.de) placed in three climate chambers at three different temperatures (ambient: 21°C, +2° 23°C and +4° 25°C). The temperature of each tank was recorded hourly over the whole experiment using *in situ* temperature loggers (HOBO Pendant MX Water Temperature Data Logger - MX2201). The tanks were equipped with LED lights (Daytime Matrix LED Pro-SLN 4000-5000K) operating on a 13:11 light:dark cycle with 3h sunrise and sundown as linear change at 8:00 am and 19:00 pm, respectively. This simulated natural daylight conditions at this specific time of the year to support phytoplankton growth and maintain ecological dynamics similar to the outdoor environment. Light intensity was set to approx. 30 µmol PAR (µmol photon m^−2^ sec^−1^).

The water for the mesocosm experiment was collected from Bunthaus station (Fig 1a, small map, including conditions on sampling day) during high tide on 29^th^ of July 2024. High tide water conditions at Bunthaus station usually reflect a phytoplankton rich water body that will be transported downstream (Hammrich 2024). We used this water body to capture water that will undergo drastic community changes observed along the Elbe Estuary (Steidle and Vennell 2024; Martens, Biederbick, and Schaum 2024). For the water collection, ten 1000 L IBC-water tanks were filled with Elbe water using a fire hose connected to a ZUWA Supersand Niro 150 pump. The water inlet for the pump was positioned at ca. 1m water depth and pumped up to 100 L per minute. The filling of the IBC tanks took ca. 90 minutes. During high tide, water body changes at the station were minimal (https://gezeiten.bsh.de/hamburg_bunthaus).

**Fig. 1:**
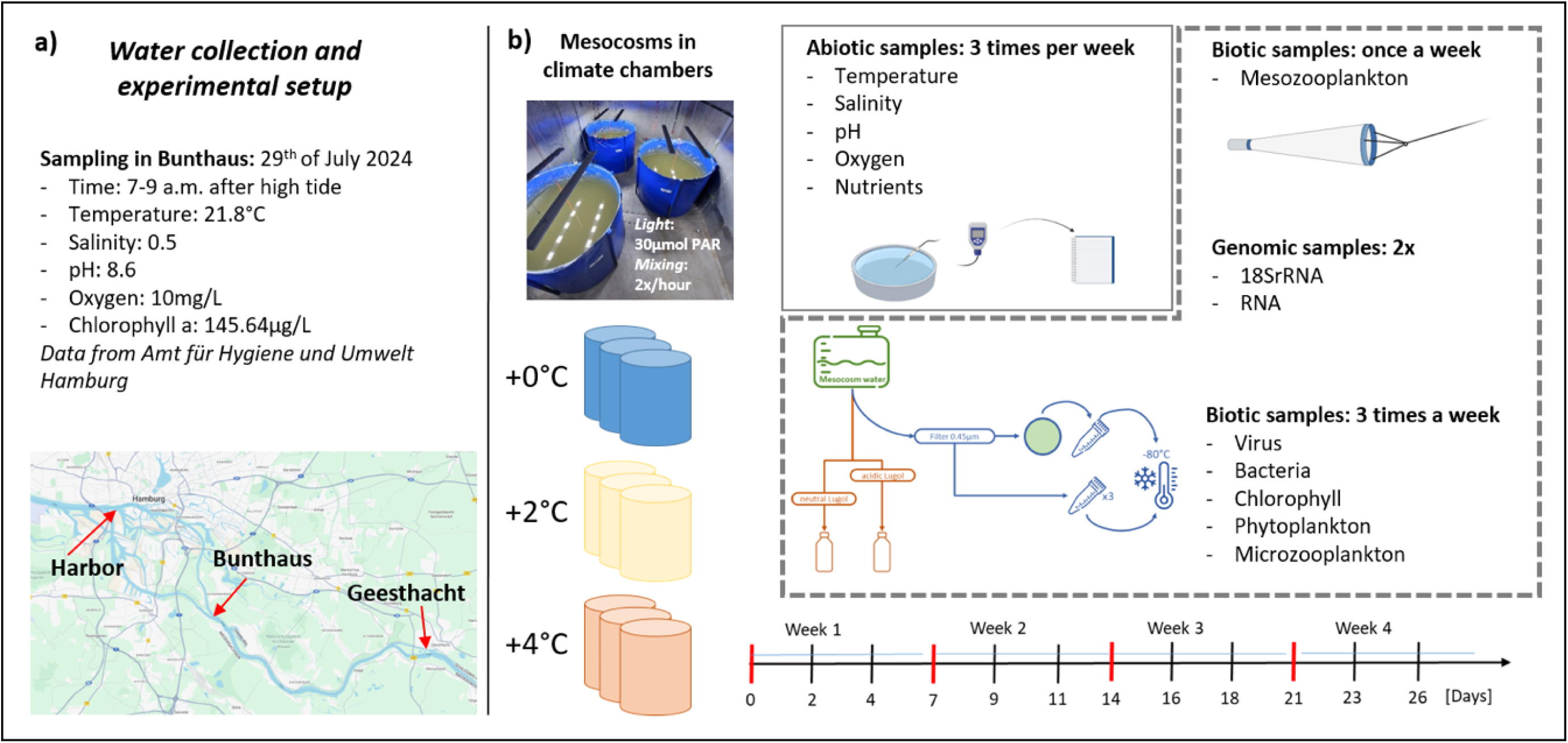
Panel a shows the time and location of water collection from river Elbe. Panel b shows the set up of the mesocosms in the climate chambers (control blue, +2°C yellow, +4°C red) and how samples were collected during the experiment.

After collection, the Elbe water was transported to the indoor mesocosm facilities and distributed in parallel via hydrostatic pressure to the nine mesocosm tanks to allow an even distribution among the tanks. Before filling, the tanks were thoroughly cleaned and degassed to prevent contamination from foreign chemicals, microorganisms and nutrients. The distribution of the water to the mesocosm tanks took roughly four hours which allowed the start of the experiment in late afternoon of the sampling day. The regular sampling started on the 31^st^ of July 2024 after a temperature acclimation period of two days.

### Sampling

Regular sampling (1^st^ sampling on the 31^st^ of July 2024) was performed throughout the four-week experimental period to monitor abiotic and biotic changes in the tanks and assess plankton dynamics in response to warming. pH, salinity, temperature and dissolved O_2_ were measured every second day using a hand-held Hanna Instruments Multiparamter device (HI98494, pH/EC/DO). Each week on Monday, Wednesday and Friday, samples for nutrient concentration, chlorophyll a (chl a) concentration, phytoplankton, microzooplankton biomass were taken. Mesozooplankton was sampled once a week. The amount of samples as well as preservation procedures are summarized in Table 1.

### Sample analysis

#### Nutrients

All nutrient analyses (nitrate, nitrite, ammonia, phosphate and silicate) followed the protocols developed by Grasshoff et al. (1983) and nutrient concentrations were determined photometrically (Spectramax id3). The samples were measured undiluted and small volumes between 1 mL to 2 mL were analyzed in triplicates.

Chlorophyll a:

Chlorophyll *a* concentrations were quantified following the protocol established by Jeffrey and Humphrey (1975). Specifically, pigments were extracted by incubating filters in 5 mL cold acetone over night at 4°C. Filters were then removed and the sample was centrifuged for 10 min. at a lot of 5000 rpm to remove cell and filter debris. The dissolved chl a was determined at wavelengths of 750, 665, 650 and 630nms and the following equation was used:

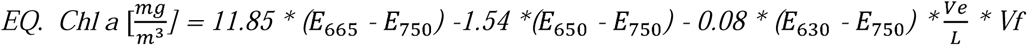

*E=extinction value, Ve= Volume of extinction, L=Path length of cuvette (cm) and Vf= dilution factor*

#### Microscopy enumeration for microphyto-, microzoo-, and mesozooplankton

Microphytoplankton and microzooplankton (<200 µm) were sampled three times per week, roughly from the middle of each tank. Samples were fixed with 0.5% neutral Lugol’s iodine solution for phytoplankton, and 1% acidic Lugol iodine solution for microzooplankton, and stored cold in darkness until analyses. For enumeration, the Utermöhl technique was used (Utermöhl 1958). Samples were transferred to Utermöhl settling chambers (10 mL chambers for phytoplankton, 25-50 mL for microzooplankton) and analysed at 200x magnification with an inverted microscope (Zeiss Axiovert 5). Either one strip of each sample or several fields of vision were counted for phytoplankton until counts of the most abundant taxa reached >400 cells (only for fields of vision). Microzooplankton was enumerated by counting either the entire or half of the bottom plate.

Mesozooplankton (>200 µm) was sampled once a week to avoid over-exploitation of the standing stocks, fixed with 4% formaldehyde and stored at room temperature. For mesozooplankton enumeration, samples were scanned using ZooSCAN (Hydroptic, CNRS patent). For the scanning procedure, dense samples were split (motoda splitter) to different ratios to avoid crowding in the scanning chamber. Scans were 8-bit greyscale, 2400 DPI images (tagged image file format; *.tif). The raw images from each scan were processed using the ZooProcess (Gorsky et al. 2010) plug-in for the program ImageJ ®, which resulted in vignettes with parameters (e.g. area, length, width, grey-level, ferret-diameter, etc.) and validated using Ecotaxa (Gorsky et al. 2010). Images per sample were summed up to determine the bulk abundance.

#### Metabarcoding analysis

On the day of water collection from Elbe river (29th of July), four samples (“field”) of 500 ml of natural Elbe water were simultaneously taken from the same location and filtered right after arriving to the mesocosm facility. After two days and three weeks of the experiment, 500 mL of water was collected in duplicates (unless otherwise stated) from each mesocosm tank. All samples were filtered through 5 µm Durapore filter to obtain the particle-associated microbial fraction (>5 µm). The filtrate was then filtered through 0.22 µm Durapore filter to obtain the free-living microbial fraction (<5 µm). After emptying the tanks at the end of the experiment the sediment was collected and preserved in 99% EtOH. All samples were immediately frozen at −80 °C. DNA and RNA from collected filters were isolated using the method described in Nercessian et al. (2005). In the samples dedicated to transcriptomic sequencing, DNA was digested with the TURBO DNA-free™ Kit (Thermo Fisher Scientific) and successful digestion was confirmed by PCR with 341F□803R primers amplifying 16S rDNA. Thereafter, cDNA synthesis was performed with ArrayScript (Thermo Fisher Scientific) with random hexamers as primers. The 18S region V9 of rDNA or rRNA was amplified using primers Euk1391F/EukBr for sequencing on the Illumina MiSeq V3 Platform. Sequencing data was analysed with QIIME2 (Bolyen et al. 2019) platform based on P2R database and further processed with the following R packages: phyloseq (McMurdie and Holmes 2013), vegan (Dixon 2003), tidyr and dplyr (Wickham et al. 2025). Visualisations were done with ggplot2 R package. The “adonis2” function from the vegan package and the “pairwiseadonis” function from pairwiseadonis R package (Martinez 2009) were applied to perform PERMANOVA and post hoc pairwise PERMANOVA statistical tests.

### Statistical analysis

All data (see previous paragraph for genomic data) were analysed using GLS (generalized least square) models (R package nlme) accounting for auto-correlation due to repeated sampling of the same tank over time and therefore controlling for tanks as random effects on the data (e.g. gls (“DATA” ∼ Temperature + Time, correlation = corAR1(form = ∼Time | tank). The GLS results are summarized in Table S1 and interpreted the following way: Differences and standard errors of the treatments (Plus 2 and Plus 4) were compared to the control intercept and if they did not overlap, we considered the treatments to have a significant effect on the analyzed data. The effect of experimental time was assessed as well and considered as a fixed factor in the model. Analysis and plotting was done in R Studio (version 4.4.3) using the following packages’: ‘ggplot2’, ‘reshape2’, ‘zoo’, ‘nlme’.

## Results

### Abiotic parameters

After the first four days of the experiment, the temperatures (Fig. 2a) remained relatively stable in each of the mesocosms (Table S1: Temperature after 200 hours; *Time* value=-0.001 +/− 0.000). There were minor temperature differences (<1°C) between the tanks due to the position of the single tanks in climate chambers, but these differences were smaller than the temperature treatment differences (Table S1: Temperature after 200 hours, *Plus 2* value=1.773+/− 0.263 and *Plus 4* value=3.840+/− 0.263). PH showed no significant differences in the temperature treatments but changed throughout the course of the experiment. Specifically, pH decreased in the first week of the experiment from 7.4 to ca. 7.0 for the control (*in situ* temperature) and +4°C treatment and 6.6 for the +2°C treatment (Fig. 2b). Then pH increased again to ca. 7.2 in all treatments (Table S1, pH, *Time* value=0.022 +/− 0.003). After a continuous increase from 0.66 to a maximum of 0.74 throughout the experiment (Table S1, Salinity, *Time* value=0.003 +/− 0.000), salinity was slightly higher in the +4°C treatment (Table S1, Salinity, Plus 4 value=0.020 +/− 0.003) at the end of the experiment (Fig. 2c).

**Fig. 2:**
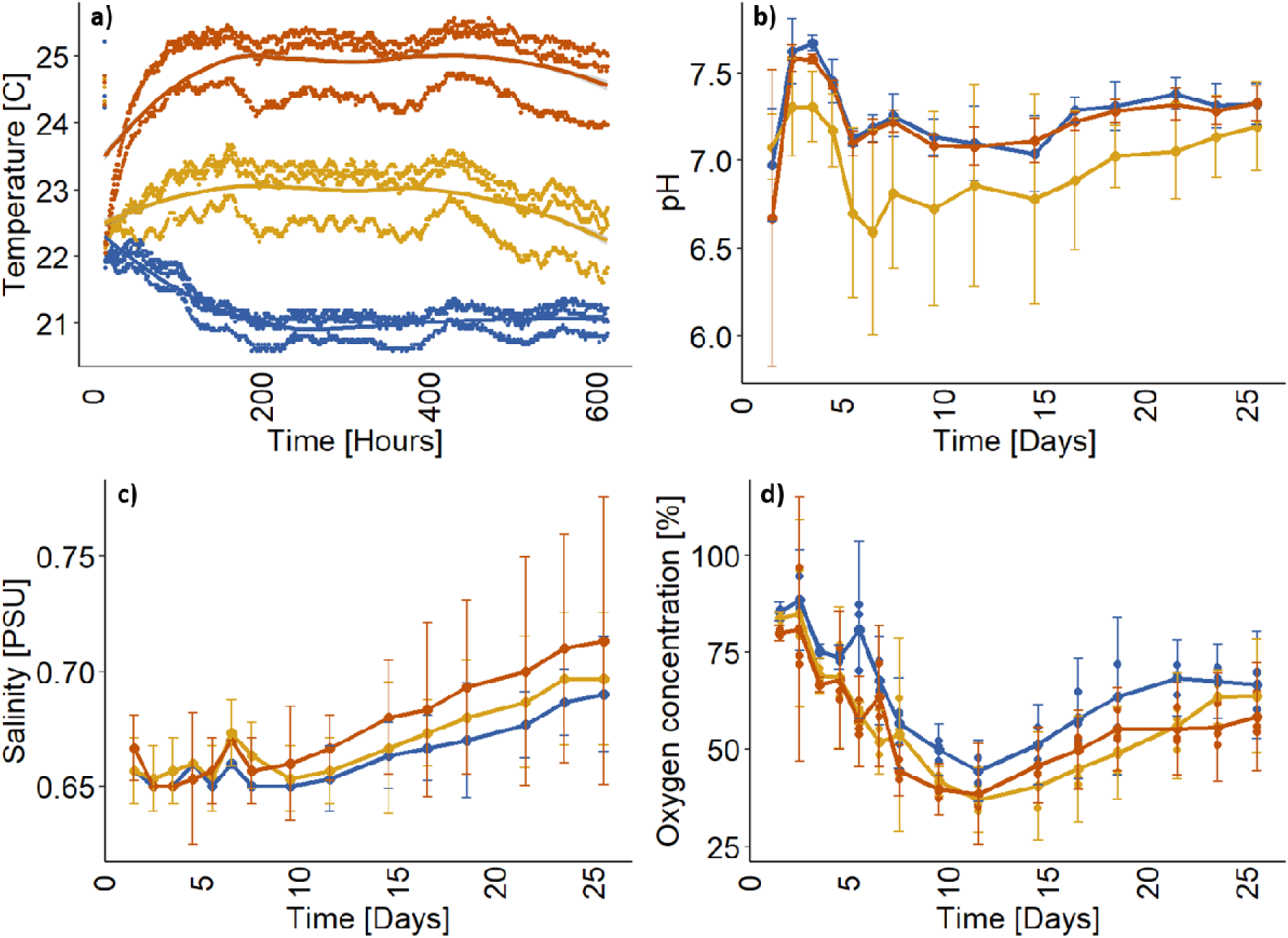
Abiotic conditions over the course of the mesocosm experiment: Colors indicate in all panels the three different experimental treatments (blue=+0°C (control), yellow=+2°C, red=+4°C). Lines represent mean of data and error bars 95% confidence intervals. In the top panels, temperature (a) and pH (b) of all nine tanks are shown. The bottom panels depict salinity (c) and oxygen concentration in % (d).

The most pronounced change in abiotic conditions was observed in oxygen concentrations (Table S1, Oxygen, *Time* value=1.705 +/− 0.140) (Fig. 2d) with ca. 8 mg mL^−1^ (90% saturation) initially and adecrease over the first 12 days to as low as 3 mg mL^−1^ (40%). Thereafter, the oxygen concentrations increased again, but did not reach initial levels and reaching a plateau at ca. 4.5-6 mg mL^−1^ (50-60%). The warming treatments showed stronger decreases in oxygen (Table S1, *Plus 2* value=-9.300+/− 1.594 and *Plus 4* value=-8.714+/− 1.594) compared to the control, indicating higher respiration with warming conditions. Note that the statistical analysis identifies a significant change over the course of the whole experiment in oxygen concentrations and between treatments (Table S1).

Temperature effects on nutrient concentrations could only be detected for ammonia but all nutrient concentrations, except for phosphate (Fig. 3 bottom left), increased slightly over time. Phosphate was just over the detection limit at ca. 0.01-0.2 µmol L^−1^ throughout the experiment. Ammonium constituted the highest nitrogen component towards the end of the experiment reaching up to 50 µmol L^−1^ (Table S1, NH4+, *Time* value=0.757 +/− 0.212). In the +4°C and control treatment, highest concentrations were reached after eight days whereas in the +2°C treatment, NH4+ increased continuously to ca. 45 µmol L^−1^at the end (Table S1, NH4+, *Plus 2* value=-16.744+/− 3.826 and *Plus 4* value=-1.207+/− 3.826) (Fig. 3 Top right). NOx slightly increased over time in the control and +2°C treatment and stayed stable in the +4°C treatment, however, overall there were no significant differences between treatments (Table S1, NOx, *Time* value=0.194 +/− 0.032) (Fig. 3 top left). Silicate increased in the second week of the experiment reaching concentrations of up to 15 µmol L^−1^ (Table S1, SiO3, *Time* value=0.559 +/− 0.037) (Fig. 3 bottom right).

**Fig. 3:**
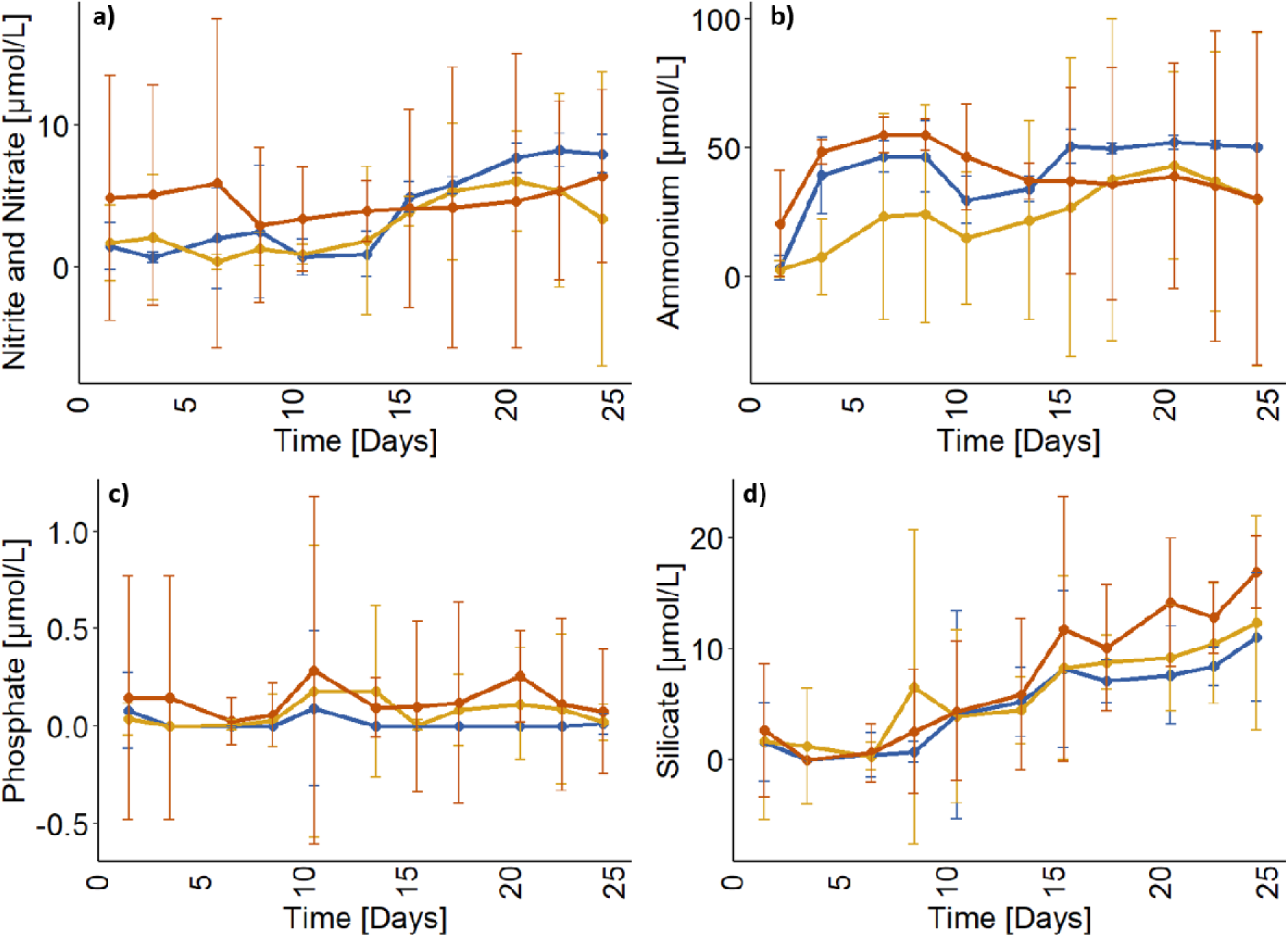
Nutrient changes over the course of the mesocosm experiment (days): Colors indicate in all panels the three different experimental treatments (blue=control, yellow=+2°C, red=+4°C). Lines represent mean of data and error bars 95% confidence intervals. In the top panels total NOx (a) and ammonia (b) of all nine tanks are shown. The bottom panels depict phosphate (c) and silicate concentrations (d).

### Plankton community changes

Similar to the abiotic changes, no strong temperature effects on the Elbe plankton community were observed (Table S1). However, autotrophs (i.e. Chl a and microphytoplankton) and heterotrophs (micro- and mesozooplankton), behaved differently. Chl *a* (Chl a)- as a proxy of autotroph bio- mass - decreased over the first ten days about 10 times (Table S1, Chl a, *Time* value=-3.816 +/− 0.415). Thereafter, a recovery of chl *a* was observed, which was highest in the +2°C treatments, but the difference between the warmed treatments was not significant (Fig. 4a). Comparable to chl *a*, the abundance of microphytoplankton (Table S1, Microphytoplankton, *Time* value=-125.469 +/− 30.250) (Fig. 4b) and microzooplankton (Table S1, Microzooplankton, *Time* value=-6844.360 +/− 948.103) (Fig. 4c) decreased in the first half of the experiment. Microphytoplankton abundance slightly recovered in the second half of the experiment but only to about 20% of the initial abundance. The recovery of microphytoplankton was highest in the control (Table S1, Microphytoplankton, *Plus 2* value=-1610.321+/− 545.668 and *Plus 4* value=-2260.252+/− 545.668) with no difference in both warmed treatments. Microzooplankton did not increase again until the end of the experiment. While chl *a*, phytoplankton and microzooplankton decreased in the first half of the experiment, mesozooplankton abundance increased over time, reaching abundances twice as high compared to initial abundances (Table S1, Mesozooplankton, *Time* value=2.990 +/− 0.436) (Fig. 4d). These changes were unaffected by temperature.

**Fig. 4.**
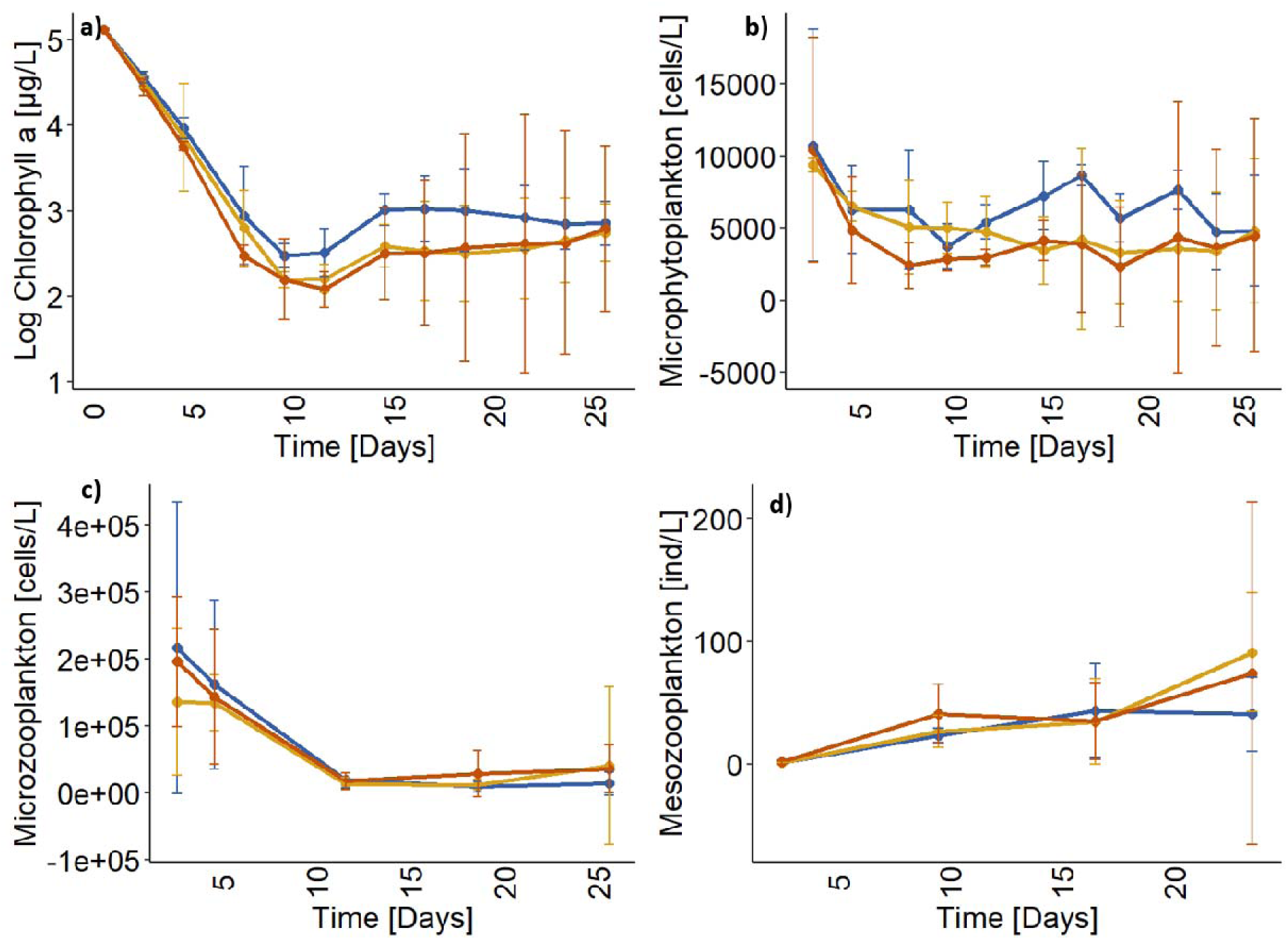
Succession of the different food-web components of the plankton over the course of the mesocosm experiment (days): Colors indicate in all panels the three different experimental treatments (blue=control, yellow=+2°C, red=+4°C). Lines represent mean of data and error bars 95% confidence intervals. In the top panels chlorophyll *a* (a) and microphytoplankton abundance (b) of all nine tanks are shown. The bottom panels depict microzooplankton (c) and mesozooplankton abundance (d).

The rDNA analysis of both size fractions (> and < 5 µm) revealed a temperature difference after three weeks of the experiment (Table 3, water 3^rd^ week). Pairwise PERMANOVA showed that these differences between size fractions and treatments were significant. In the Principal Coordinate Analysis (PCA), the first PC axis best explained the difference between the size fractions (Fig. 5a, closed and filled circles and squares), whereas the second PC axis best explained differences between sampling days and thus over time (Fig. 5a, circles and squares). This indicates that the community shifted over time and in response to temperature. While PERMANOVA tests were not significant for rRNA data, the ordination plot revealed two things: first the size fractions each clustered together but are different from each other (Fig. 5b, filled and closed squares). Second, there was a “mesocosm effect” at the transcriptome level, especially in the smaller size fraction (Fig 5b green squares vs. rest, Table 2). Furthermore, there was no significant temperature effect at the rRNA transcriptome levels right after acclimatization (Table 2.). The sediment samples taken after three weeks of the experiment clustered with the water samples of the >5 µm size fraction taken during the first week of the experiment (Fig. 5a) indicating similarities to the community found in the water column at the beginning of the experiment.

**Fig. 5.**
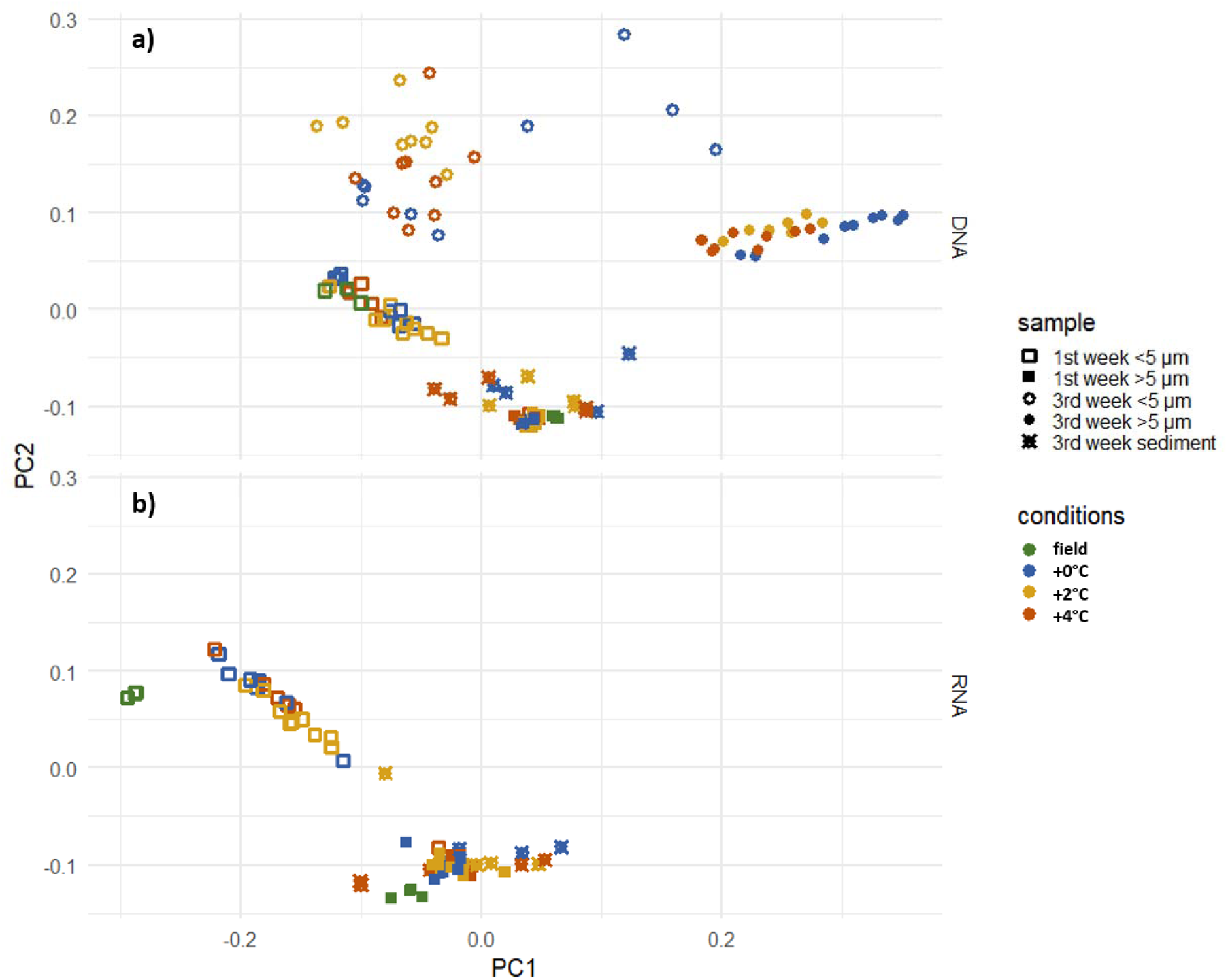
Principal Coordinate Analysis of 18S rDNA (panel a) and rRNA (panel b) between two sampling times - first week and third week. Colors indicate the three different experimental treatments (blue=control, yellow=+2°C, red=+4°C). Signs shapes indicate the time of sampling and sample type: sediment or water. Filled or empty shapes differentiate two water sample size fractions: >5 µm and <5 µm.

**Table 2.** PERMANOVA test results across time of experiment. Significant test results (P-value ≤ 0.05) are marked with (*).

| Sample type | Size | Week | Conditions | Data | F | R2 | P-value |
| --- | --- | --- | --- | --- | --- | --- | --- |
| Water | >5 µm | 1st week | mesocosm vs field | rDNA | 0.644 | 0.027 | 0.602 |
| Water | <5 µm | 1st week | mesocosm vs field | rDNA | 1.398 | 0.057 | 0.182 |
| Water | <5 µm | 1st week | mesocosm vs field | rRNA | 0.614 | 0.026 | 0.592 |
| Water | >5 µm | 1st week | mesocosm vs field | rRNA | 0.931 | 0.039 | 0.402 |
| Water | >5 µm | 1st week | temperatures | rDNA | 0.837 | 0.085 | 0.536 |
| Water | < 5µm | 1st week | temperatures | rDNA | 0.725 | 0.071 | 0.664 |
| Water | >5 µm | 1st week | temperatures | rRNA | 1.823 | 0.168 | 0.073 |
| Water | <5 µm | 1st week | temperatures | rRNA | 1.32 | 0.122 | 0.244 |
| Water | >5 µm | 3rd week | temperatures | rDNA | 4.770 | 0.389 | 0.001 (*) |
| Water | < 5µm | 3rd week | temperatures | rDNA | 3.976 | 0.346 | 0.001 (*) |
| Sediment | - | 3rd week | temperatures | rDNA | 0.785 | 0.136 | 0.604 |
| Sediment | - | 3rd week | temperatures | rRNA | 0.785 | 0.136 | 0.577 |

**Table 3.** Post-hoc pairwise PERMANOVA results for rDNA data after three weeks of experiment of each size fraction and all temperatures comparisons. Significant test results (P-value ≤ 0.05) are marked with (*).

| Size fraction + conditions | F | R2 | P-value |
| --- | --- | --- | --- |
| >5 $\mu\text{m}$ +2°C vs. control | 5.265 | 0.345 | 0.011 (*) |
| >5 $\mu\text{m}$ +2°C vs. +4°C | 1.485 | 0.129 | 0.192 |
| >5 $\mu\text{m}$ +4°C vs. control | 7.493 | 0.428 | 0.006 (*) |
| <5 $\mu\text{m}$ +2°C vs. control | 5.211 | 0.343 | 0.005 (*) |
| <5 $\mu\text{m}$ +2°C vs. +4°C | 2.241 | 0.183 | 0.05 (*) |
| <5 $\mu\text{m}$ +4°C vs. control | 4.279 | 0.300 | 0.007 (*) |

## Discussion

We here present the first experimental mesocosm study investigating the Elbe River plankton community and explore how warming affects abiotic conditions and subsequently plankton community. With this study we also show, how an experimental mesocosm setup can shed light on complex riverine dynamics despite its controlled and semi-natural setup that can also influence experimental outcomes. Our four-week summer mesocosm experiment revealed strong phenological shifts in the plankton community with potential trophic interactions due to differing responses on the trophic levels. Temperature driven community changes can have cascading effects on biogeochemical and ecological processes in an estuarine system. However, warming usually does not function as a single stressor but in concert with other (anthropogenic) stressors, i.e. pollution that can sometimes hide or override patterns like strong overall temporal changes in the plankton community and abiotic parameters.

### Warming effects on abiotic conditions

The warming-induced changes in the abiotic parameters during the experiment provide insight into the ongoing biogeochemical processes within the lower food. Especially prominent was the strong decline in oxygen to less than 50% saturation, which was even more intense in the warming treatments, in the first half of the experiment followed by recovery thereafter, which was higher at ambient temperatures (control). The strong and uniform decline, likely resulted from intensified heterotroph respiration associated with re-mineralization of organic matter and elevated metabolic activity under stressful conditions (Falkowski, Barber, and Smetacek 1998; von Scheibner et al. 2014) which are known to be strong in “artificial” systems and experimental setups (Stewart et al. 2013). Oxygen depletions are a recurrent phenomenon in the Elbe River (Schöl et al. 2014) and while mesocosms represent semi-natural and enclosed systems (Stewart et al. 2013) that have their own intricate dynamics, we compared the changes in oxygen concentrations in the mesocosms to observed oxygen dynamics in the Elbe Estuary at the same time. At the end of July and beginning of August 2024, there was a drastic oxygen depletion along the lower part of the Elbe Estuary (personal communication Hammrich, A. 2025) where oxygen decreased to as low as 2 mg L^−1^ in the Hamburg harbour area and increased again up to 6 mg L^−1^ further downstream. Based on modeling data (depending on tides and river run-off) (Hammrich 2024) water usually travels for a period of 40-50 days from Geesthacht to Brunsbüttel (river mouth) and would therefore take roughly 6-7 days from Bunthaus to the harbor (the stretch with the strongest changes in oxygen) (Fig. 1, Map). The striking similarity in absolute oxygen concentrations between temporal oxygen dynamics in the mesocosms and spatial oxygen gradients in the river system suggests that water traveling through the harbor system may become partially isolated, allowing processes that drive substantial oxygen declines. Changes in plankton community composition may play an important role in explaining these patterns and warrant further investigation.

The targeted temperature was reached in the respective mesocosms after three days and remained relatively stable throughout the experiment, thus, successfully mimicking a warming (heat-wave) event in the Elbe in summer. The slight salinity increase, more pronounced in the warmest treatment, can likely be attributed to evaporation effects, which were stronger in the +4°C treatment with highest water loss. Concurrent with oxygen dynamics, the pH decrease observed within the first week could result from shifts in carbon chemistry driven by increased microbial respiration. Elevated respiration produces CO□, which dissolves into the water and forms carbonic acid, thereby acidifying the water (Jin and Liu 2023). While phytoplankton influence pH through photosynthesis resulting in increases in pH at high photosynthetic activity (López-Urrutia et al. 2006), the rapid initial decline in pH in this experiment suggests that microbial respiration played a predominant role in pH fluctuations during the early phase. Directly measuring photosynthesis and respiration could help to understand how primary producers and heterotrophs react differently to the stressful conditions in the beginning of the experiment and later to warming.

Even though changes in nutrient dynamics in our system were small and the start of the experiment coincided with a low nutrient season (Kamjunke and Sanders 2024), the dynamics highlight the processes occurring within the studied system. The observed rise in NOx concentrations (NO□+ NO□) following an increase in ammonia indicates active remineralization of organic nitrogen through nitrification pathways (Sanders, Schöl, and Dähnke 2017). Organic material could stem from sedimentation (indicated by the clustering of sediment and initial community samples in our genomic analysis) of decaying plankton or grazing activity by micro- and mesozooplankton in the system that produce fecal pellets (Tang et al. 2006) and detritus by sloppy feeding (Steinberg and Landry 2016). The initial ammonium surge, followed by NOx increase, suggests microbe mediated conversion of organic nitrogen to inorganic forms, with NO□ likely representing the primary component of NOx in this setting (Kamjunke et al. 2023). In future mesocosm studies, identification of the bacterial community composition would shed light on the potential presence of denitrifying bacteria. Phosphate levels remained low throughout the experiment, reflecting the general phosphate limitation characteristic of the Elbe River in summer (Hardenbicker et al. 2016). In contrast, silicate concentrations increased notably during the second part of the experiment, implying silicate release during phytoplankton loss resulting from cell senescence and grazing activity by micro- and mesozooplankton (Steinberg and Landry 2016). For active remineralization of biogenic silica from diatom frustules or other siliceous organisms, the experimental time of three weeks was most likely not long enough to achieve already an advanced status of remineralization (Bidle and Azam 1999; Kamjunke et al. 2023).

### Warming effects on the plankton community

Primary producers (measured as chl *a* and microphytoplankton abundances) declined substantially within the first ten days, paralleling the decrease in oxygen. The initial reduction happened most likely due to high grazing pressure by mesozooplankton and the sink-out of large-sized microphytoplankton due to reduced mixing of the water column in our experimental set-up. Further, microzooplankton that is considered a principal grazer of phytoplankton (Landry and Calbet 2004) likely contributed to the observed phytoplankton decrease in addition. However, definitive mechanistic links require further analyses. Similar to microphytoplankton, microzooplankton abundances declined sharply during the early phase of the experiment, although a minimum was reached two days after that of the microphytoplankton. The decline in microphytoplankton and microzooplankton was most likely related to an intense top-down control by mesozooplankton that showed a strong increase in abundances in the same period. In general, feeding preferences of mesozooplankton depend strongly on the abundance and taxonomic composition of phytoplankton and microzooplankton and on feeding modes of mesozooplankton (Sommer, Hansen, and Sommer 2006). Further, the nutrient status in the water column plays a pivotal role by altering the nutritional quality of phytoplankton while microzooplankton is considered to buffer nutritional imbalances efficiently thus resulting in trophic upgrading and increasing the prey quality for consumers (Klein Breteler et al. 1999; Malzahn et al. 2010; Šorf et al. 2014). Under nutrient-limited conditions like in the Elbe River in summer, a strong predation of mesozooplankton on microzooplankton prey can thus be expected since microzooplankton is considered as a high-quality prey item compared to phytoplankton when nutrients are limited (Löder et al. 2011).

### Community analyses using metabarcoding

Genomic monitoring is increasingly recognized as essential for understanding how species and communities respond to rapid climate change through shifts in gene expression, adaptive variation, and community composition (Bernatchez et al. 2023). By linking changes in gene expression (in essence RNA analysis), taxonomic signatures, or functional gene abundances to experimentally measured alterations in productivity, nutrient cycling, and trophic interactions, we gain a more mechanistic and integrative understanding of how temperature influences ecosystem processes.

This further strengthens experimental interpretations and improves predictions of estuarine plankton responses to warming. No significant differences between both rDNA and rRNA for different temperatures at the start of the experiment confirm equal starting conditions. A temperature effect on rDNA sequences was only detectable in the second half of the experiment, most likely due to hidden patterns caused by strong shifts within the plankton community in the beginning. The grouping of sediment samples with those samples of > 5µm in the water column taken during the first week of the experiment suggests the sink-out of cells of this fraction that sedimented to the bottom of the mesocosm tanks.

### Summary and conclusion

The relatively subtle temperature effects observed in our mesocosms may arise from a combination of biological and experimental factors, which has also been documented in similar studies. First, natural plankton communities often show high context-dependence and community composition dynamics that can mediate or dampen direct thermal responses. This is because trophic interactions and community structure can change more strongly than baseline temperature sensitivities such as warming effects on specific functional groups, which vary with season and community composition (Soulié et al. 2022). Second, mesocosm systems can generate strong internal dynamics, such as trophic cascades, oxygen depletion, and shifts in grazing pressure, which may override or mask temperature effects on individual metabolic rates. This can be especially prominent over short experimental periods, which was, e.g., shown for phytoplankton biomass dynamics being driven more by grazing than by temperature alone in a Baltic Sea mesocosm study (Paul et al. 2016). Third, the magnitude and duration of warming treatments relative to natural variability and acclimation potential of organisms can constrain detectable responses. Different species and taxa exhibit differences in temperature optima and tolerances as well as acclimation and compensatory responses that reduce apparent temperature sensitivity over ecologically relevant timescales (organisms with metabolic plasticity can buffer ecosystem-level responses to moderate warming) (Kordas et al. 2022).

Despite certain experimental limitations, mesocosm experiments remain powerful tools for climate change ecology because they integrate multiple interacting drivers under semi-natural conditions while allowing controlled manipulations (Grossart et al. 2026). Mesocosm experiments capture complex species interactions, community reassembly, and joint responses of physical, chemical, and biological processes that cannot be reproduced in single-species laboratory assays (Sommer et al. 2007b; Stewart et al. 2013; Lewandowska et al. 2014). For example, warming in larger multitrophic mesocosms can alter plankton body size spectra and trophic interactions in ways that inform both mechanistic understanding and model development (Albini et al. 2025; Stewart et al. 2013). Additionally, by at least initially maintaining natural diversity and allowing continuous monitoring of functional processes (e.g., oxygen dynamics, primary production, metabolic coupling), mesocosms can bridge the gap between controlled experimentation and field realism, which is essential for predicting ecosystem-scale responses to future climate scenarios.

## Acknowledgements

We would like to thank our student helpers and lab technicians of IMF for their help and support in this study. In addition we would like to thank Werner Blohm and Michael Lechelt of the “Amt für Hygiene und Umwelt” of the city of Hamburg for providing their facilities for sampling. ChatGPT (OpenAI, GPT-5.1) was used for language and grammar improvement, and all outputs were reviewed and validated by the authors.

## Author contribution

- Project conceptualization: LL, NA, AMM
- Development of experimental setup: LL, ML, DNP, SP, NA, AMM
- Running of Experiment and sample analysis: LL, ML, DNP, SP
- Data analysis: LL, JG
- Manuscript writing: LL
- Manuscript editing: NA, AMM, JG, HPG, ML, ES, DNP, SP
- Funding acquisition: LL, NA, AMM, ES, HPG
- All authors agreed to last version of the manuscript

## Data availability

The data and analysis for abiotic and biotic measurements are submitted and available: 10.5281/zenodo.18348948

The sequencing data (OTUs) and analysis are available: https://github.com/JuliaGol/Elbe_mesocosms_warming_experiment

